# Genome-wide local ancestry and direct evidence for mitonuclear coadaptation in African hybrid cattle populations (*Bos taurus*/*indicus*)

**DOI:** 10.1101/2021.08.26.457829

**Authors:** James A. Ward, Gillian P. McHugo, Michael J. Dover, Thomas J. Hall, Said Ismael Ng’ang’a, Tad S. Sonstegard, Daniel G. Bradley, Laurent A. F. Frantz, Michael Salter-Townshend, David E. MacHugh

## Abstract

The phenotypic diversity of African cattle reflects adaptation to a wide range of agroecological conditions, human-mediated selection preferences, and complex patterns of admixture between the humpless *Bos taurus* (taurine) and humped *Bos indicus* (zebu) subspecies, which diverged 150-500 thousand years ago. Despite extensive admixture, all African cattle possess taurine mitochondrial haplotypes, even populations with significant zebu biparental and male uniparental nuclear ancestry. This has been interpreted as the result of a human-mediated dispersal ultimately stemming from zebu bulls imported from South Asia during the last three millennia. Here we assess whether ancestry at mitochondrially-targeted nuclear genes in African admixed cattle is impacted by mitonuclear functional interactions. Using high-density SNP data, we find evidence for mitonuclear coevolution across hybrid African cattle populations with a significant increase of taurine ancestry at mitochondrially-targeted nuclear genes. Our results, therefore, support the hypothesis of incompatibility between the taurine mitochondrial genome and the zebu nuclear genome.

**GRAPHICAL SUMMARY:** 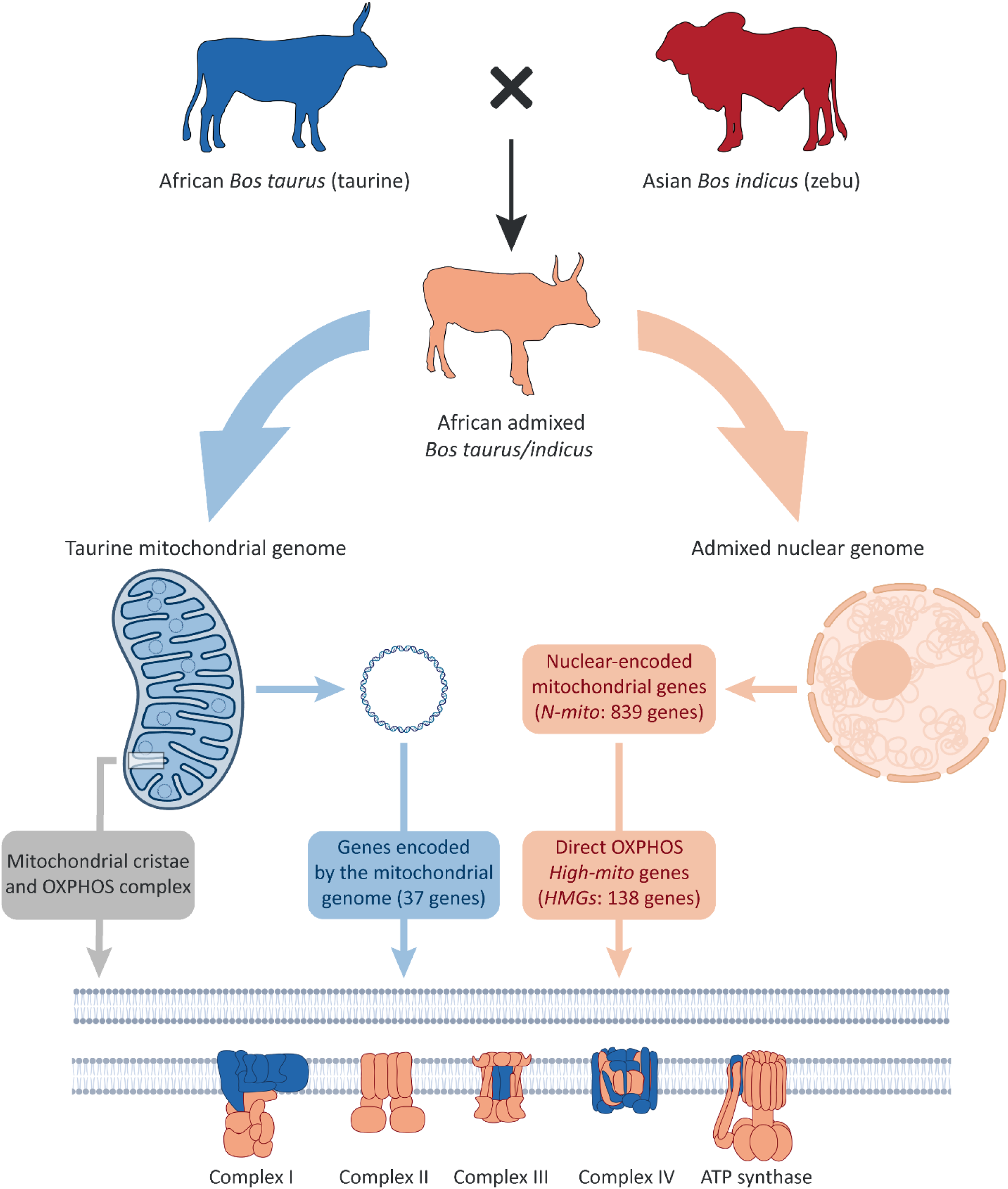

**Highlights:**

• Using high-density genome-wide SNP data, we present evidence for mitonuclear coevolution in hybrid African cattle.
• We observe a significant increase of taurine ancestry across multiple hybrid populations at mitochondrially-targeted nuclear genes.
• Our results provide support for the hypothesis of mitonuclear incompatibility between the zebu nuclear genome and the taurine mitochondrial genome.

## INTRODUCTION

Hybridization between divergent lineages results in an influx of new genetic variants which can improve the adaptive potential of animal and plant populations (Hedrick, 2013; Moran et al., 2021). It has long been used by breeders to generate livestock populations with specific phenotypic characteristics (Wu and Zhao, 2021). For example, crossbreeding between Asian and European domestic pigs, which diverged ∼1 million year ago, was used by 19^th^ century European breeders as a strategy to improve the fertility of local landraces (Bosse et al., 2014; White, 2011).

Human-mediated crossbreeding between humpless *Bos primigenius taurus* (*B. taurus* – taurine) and humped *Bos primigenius indicus* (*B. indicus* – zebu), which diverged 150 to 500 kya (Chen et al., 2018; Wang et al., 2018; Wu et al., 2018), has also played a major role in shaping the genetic composition of many African cattle populations. In fact, recent nuclear genome studies have shown that cattle ancestry in Africa represents a mosaic shaped by admixture between the original substrate of locally adapted taurine cattle, which likely first came to Africa with people during the Neolithic period, and more recently introduced South Asian zebu (Kim et al., 2017; Kim et al., 2020). This long process of admixture, which likely lasted thousands of years (Verdugo et al., 2019), led to the establishment of indigenous African cattle populations that are deeply rooted in rural African communities, forming an integral part of food production and cultural and religious activities throughout the continent (Van Marle-Köster et al., 2021).

Despite extensive admixture, however, all mitochondrial genomes of native African cattle populations analyzed to-date exclusively cluster within the taurine T1 haplogroup (Bradley et al., 1996; Kwon et al., 2022; Loftus et al., 1994a; Loftus et al., 1994b; Troy et al., 2001). This observation, together with the widespread distribution of *B. indicus* Y-chromosome haplotypes across Africa (Hanotte et al., 2000; Perez-Pardal et al., 2018), has been interpreted as the result of human-mediated dispersal and breeding of zebu bulls from South Asia during the last three millennia (Hanotte et al., 2002; Hanotte *et al*., 2000; MacHugh et al., 1997; Perez-Pardal *et al*., 2018).

Functional mismatch between the mitochondrial and nuclear genomes transmitted from two divergent parental lineages have been observed in many vertebrate populations (Hill, 2019; Hill et al., 2019). For example, recent studies of hybridization in hares, sparrows, and hominids, have provided compelling evidence for mitonuclear incompatibilities (Seixas et al., 2018; Sharbrough et al., 2017; Trier et al., 2014). These likely stem from the fact that the 37 genes located in vertebrate mitochondrial genomes (Boore, 1999), also rely on over one thousand co-adapted nuclear genes that encode proteins and protein subunits essential to the efficient function of the mitochondrion (Blier et al., 2001; Rand et al., 2004; Sloan et al., 2018; Woodson and Chory, 2008). The most well studied example of mitonuclear cooperation is the oxidative phosphorylation (OXPHOS) system, which consists of five protein complexes, four of which are chimeric— assembled using subunits encoded both by the nuclear and mitochondrial genomes (Allen, 2015; Isaac et al., 2018; Rand *et al*., 2004). Mitonuclear incompatibilities between distinct inter- and intraspecific evolutionary lineages can give rise to deleterious biochemical effects associated with reduced efficacy of OXPHOS protein complexes (Ballard and Melvin, 2010; Blier *et al*., 2001; Ellison and Burton, 2006; Ellison et al., 2008), which leads to lower ATP production (Ellison and Burton, 2006; Ellison *et al*., 2008; McKenzie et al., 2003; McKenzie et al., 2004) and increased levels of oxidative damage (Barreto and Burton, 2013; Du et al., 2017; Latorre-Pellicer et al., 2016; Pichaud et al., 2019).

Fixation of the T1 haplogroup in African cattle has been investigated recently. An approximate Bayesian computation (ABC) approach using genome-wide nuclear SNP data from 162 East African cattle indicated that a model of male-mediated dispersal combined with mitonuclear interactions could explain current patterns of bovine genomic diversity in this region (Kwon *et al*., 2022). Here, we examine continent-wide discordance of uniparental and biparental genomic variation in African cattle and test the hypothesis that functional incompatibilities have arisen between the mitochondrial and nuclear genomes in hybrid cattle populations across the continent (Figure 1). To do this, we analyzed high-density SNP data encompassing the nuclear and mtDNA genomes (Illumina^®^ BovineHD 777K BeadChip) from 678 animals representing 18 African, Asian, and European breeds/populations) and 174 complete bovine mitochondrial genomes. These data were used to characterize genome-wide local ancestry and systematically evaluate mitonuclear interactions, coadaptation, and functional mismatch in multiple genetically independent admixed African cattle populations.

**Figure 1.**
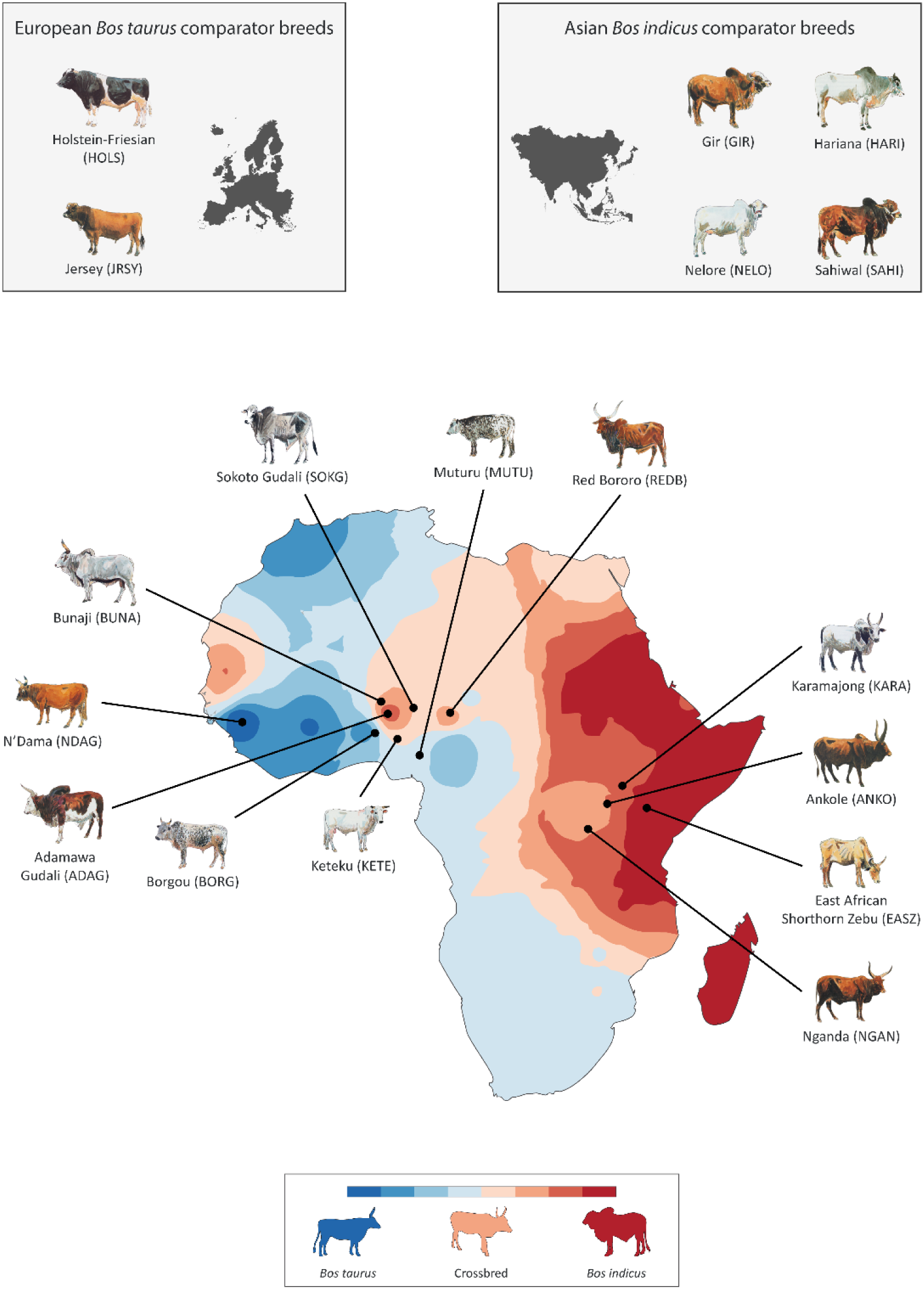
Geographical patterns of African *Bos taurus* and Asian *Bos indicus* admixture in hybrid African cattle populations. Map of Africa showing sampled cattle populations and an interpolated synthetic map illustrating spatial distribution of African *B. taurus* and Asian *B. indicus* admixture. Also shown are two European *B. taurus* and four Asian *B. indicus* comparator breeds. Admixture data was generated from the first principal component (PC1) of a principal component analysis (PCA) of microsatellite genetic variation across African cattle populations (Hanotte *et al*., 2002). Modified from (McHugo et al., 2019) under the terms of the Creative Commons Attribution 4.0 International License (http://creativecommons.org/licenses/by/4.0). (*B*) Mitonuclear interactions that can give rise to mitonuclear incompatibilities in crossbred and hybrid African cattle populations. All African cattle surveyed to-date retain the taurine mitochondrial genome (some figure components created with a <u>BioRender.com</u> license).

## RESULTS AND DISCUSSION

### Complex mitonuclear genomic structure in African admixed cattle

We first established the ancestry of the animals in our sample set using the BovineHD 777K BeadChip data. Filtering and quality control of the BovineHD 777K BeadChip resulted in 562,635 SNPs and 605 individual animals retained for subsequent analyses (Table 1). Figure 2A shows a PCA plot generated using SNP genotype data for Asian *B. indicus*, East and West African admixed *B. indicus/taurus*, African *B. taurus*, and European *B. taurus* cattle. PC1 (58.4%) and PC2 (17.9%) account for the bulk of the variance and represent the splits between *B. indicus* and *B. taurus* and the African and European taurine lineages, respectively. The results of the genetic structure analysis using the fastSTRUCTURE program and an inferred number of clusters of *K* = 3 are shown in Figure 2B, which illustrates taurine and zebu autosomal genomic ancestry across individual East and West African admixed animals and breeds (Figure S1 and Table S1). These results recapitulate, at higher resolution, continent-wide patterns of admixture that were previously observed using smaller panels of microsatellite and SNP markers (Decker et al., 2014; Hanotte *et al*., 2002).

**Figure 2.**
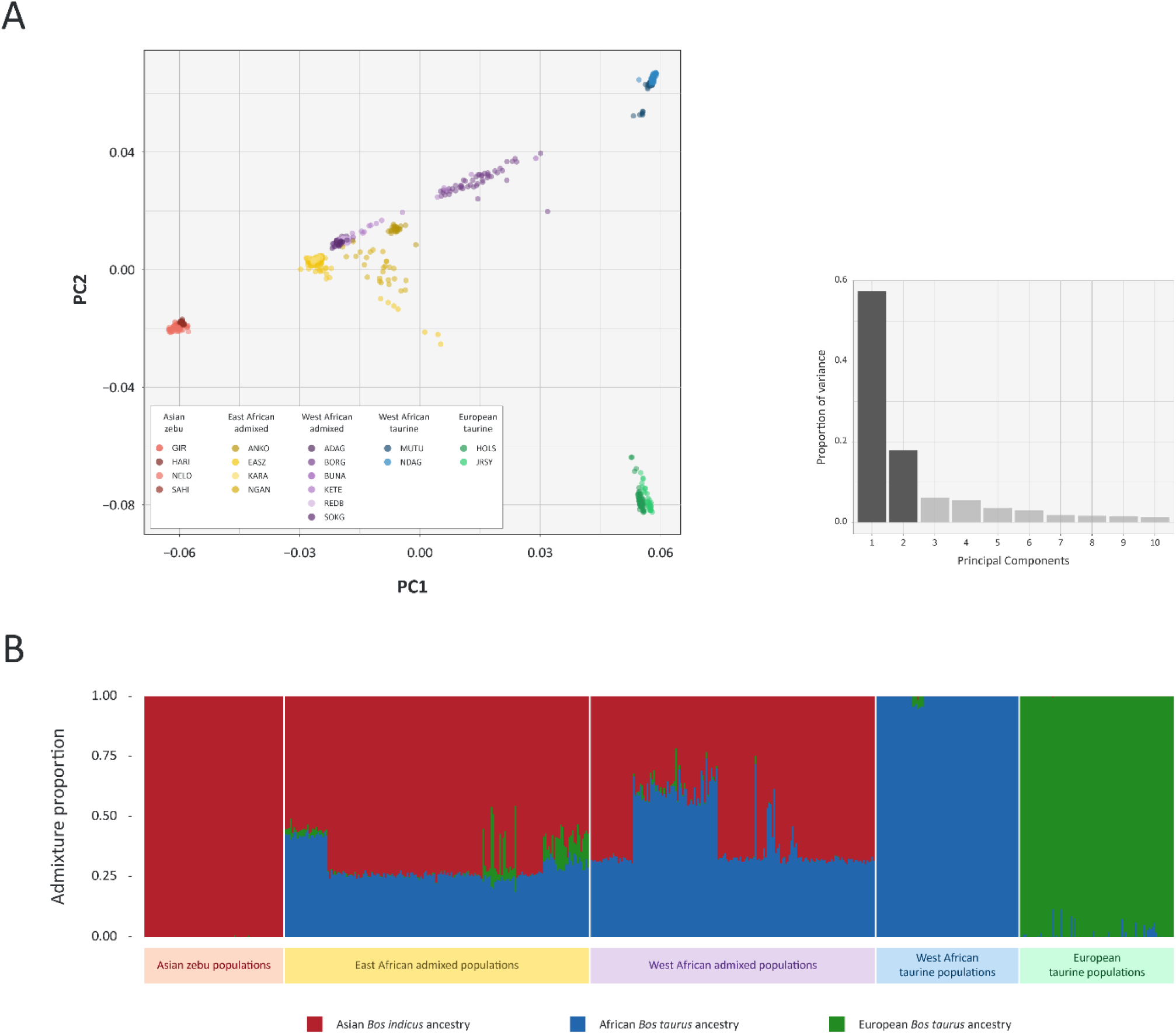
Autosomal genomic diversity and admixture in African, Asian, and European cattle. (A) Results of the principal component analysis (PCA) for 605 animals from 18 different cattle breeds genotyped for 562,635 SNPs. The PCA plot shows the coordinates for each animal based on the first two principal components. Principal component 1 (PC1) differentiates the *Bos taurus* and *Bos indicus* evolutionary lineages, while PC2 separates the African and European taurine groups. A histogram plot of the relative variance contributions for the first 10 PCs is also shown with PC1 and PC2 accounting for 58.4% and 17.9% of the total variation for PC1–10, respectively. (B) Unsupervised genetic structure plot for Asian zebu, East and West African admixed cattle, and West African and European taurine breeds. Results for an inferred number of ancestry clusters of *K* = 3 is shown, which corresponds to Asian *Bos indicus* (red), European *Bos taurus* (green), and African *B. taurus* (blue) ancestral components, respectively.

**Table 1.** Cattle breeds/populations, geographical origins, and sources of BovineHD 777K SNP data.

| Cattle breed/population | Code | Type/morphology | Country of origin | Source 1<sup>a</sup> | Source 2<sup>b</sup> | Source 3<sup>c</sup> | Source 4<sup>d</sup> | Total (n) |
| --- | --- | --- | --- | --- | --- | --- | --- | --- |
| Muturu | MUTU | West African taurine | Nigeria | 8 | --- | 20 | --- | 28 |
| N'Dama | NDAG | West African taurine | Guinea | 24 | 9 | --- | 23 | 56 |
| Holstein-Friesian | HOLS | European taurine | Netherlands | 59 | --- | --- | --- | 59 |
| Jersey | JRSY | European taurine | United Kingdom | 32 | --- | --- | --- | 32 |
| Ankole | ANKO | East African admixed | Uganda | 25 | --- | --- | --- | 25 |
| East African Shorthorn Zebu | EASZ | East African admixed | Kenya | 92 | --- | 19 | --- | 111 |
| Karamojong | KARA | East African admixed | Uganda | 16 | --- | --- | --- | 16 |
| Nganda | NGAN | East African admixed | Uganda | 23 | --- | 4 | --- | 27 |
| Adamawa Gudali | ADAG | West African admixed | Nigeria | 23 | --- | 2 | --- | 25 |
| Borgou | BORG | West African admixed | Benin | --- | --- | --- | 50 | 50 |
| Bunaji | BUNA | West African admixed | Nigeria | 22 | --- | 5 | --- | 27 |
| Keteku | KETE | West African admixed | Nigeria | --- | --- | 22 | --- | 22 |
| Red Bororo | REDB | West African admixed | Nigeria | 22 | --- | 4 | --- | 26 |
| Sokoto Gudali | SOKG | West African admixed | Nigeria | 19 | --- | --- | --- | 19 |
| Gir | GIR | Asian zebu | India | 28 | --- | --- | --- | 28 |
| Hariana | HARI | Asian zebu | India | --- | 10 | --- | --- | 10 |
| Nelore | NELO | Asian zebu | Brazil | 34 | --- | --- | --- | 34 |
| Sahiwal | SAHI | Asian zebu | India | --- | 10 | --- | --- | 10 |

<sup>a</sup> Bahbahani *et al.* (2015); <sup>b</sup> Verdugo *et al.* (2019); <sup>c</sup> Acceligen cattle genotyping database; <sup>d</sup> genotyped for the present study.

After filtering of the 346 mtDNA SNPs on the BovineHD 777K BeadChip and identification of ancestry-informative SNPs that distinguish the taurine and zebu mtDNA genomes, a haplotype network was generated using 39 mtDNA SNPs and a total of 491 cattle (47 African taurine, 82 European taurine, 156 East African admixed, 136 West African admixed, and 70 Asian zebu). Figure 3A shows this network and demonstrates that all 339 African taurine and admixed cattle surveyed here possess the taurine mitochondrial genome. In this respect, animals with predominantly zebu ancestry and morphology in Africa represent an example of “massively discordant mitochondrial introgression” (MDMI) (Bonnet et al., 2017), most likely as a result of male-mediated gene flow and genetic drift through preferential dissemination of *B. indicus* genetic material by a relatively small number of Asian zebu cattle, most of which were bulls (Bradley et al., 1994; Loftus *et al*., 1994a). This scenario is strongly supported by the widespread dissemination of the *B. indicus* Y chromosome in African admixed and morphologically taurine cattle populations (Hanotte *et al*., 2000; Perez-Pardal *et al*., 2018). In addition, taurine-zebu uniparental and biparental genomic structure on the continent has been influenced by specific livestock breeding practices, adaptation to savanna biomes, cultural preferences, and the logistics of long-distance terrestrial and maritime trade networks encompassing Southern Asia, Arabia, and North and East Africa (Boivin et al., 2014; Boivin and Fuller, 2009; Gifford-Gonzalez and Hanotte, 2011; Marshall, 1989), in conjunction with massive cattle replacements following the rinderpest panzootics of the late 19th century (Spinage, 2003).

**Figure 3.**
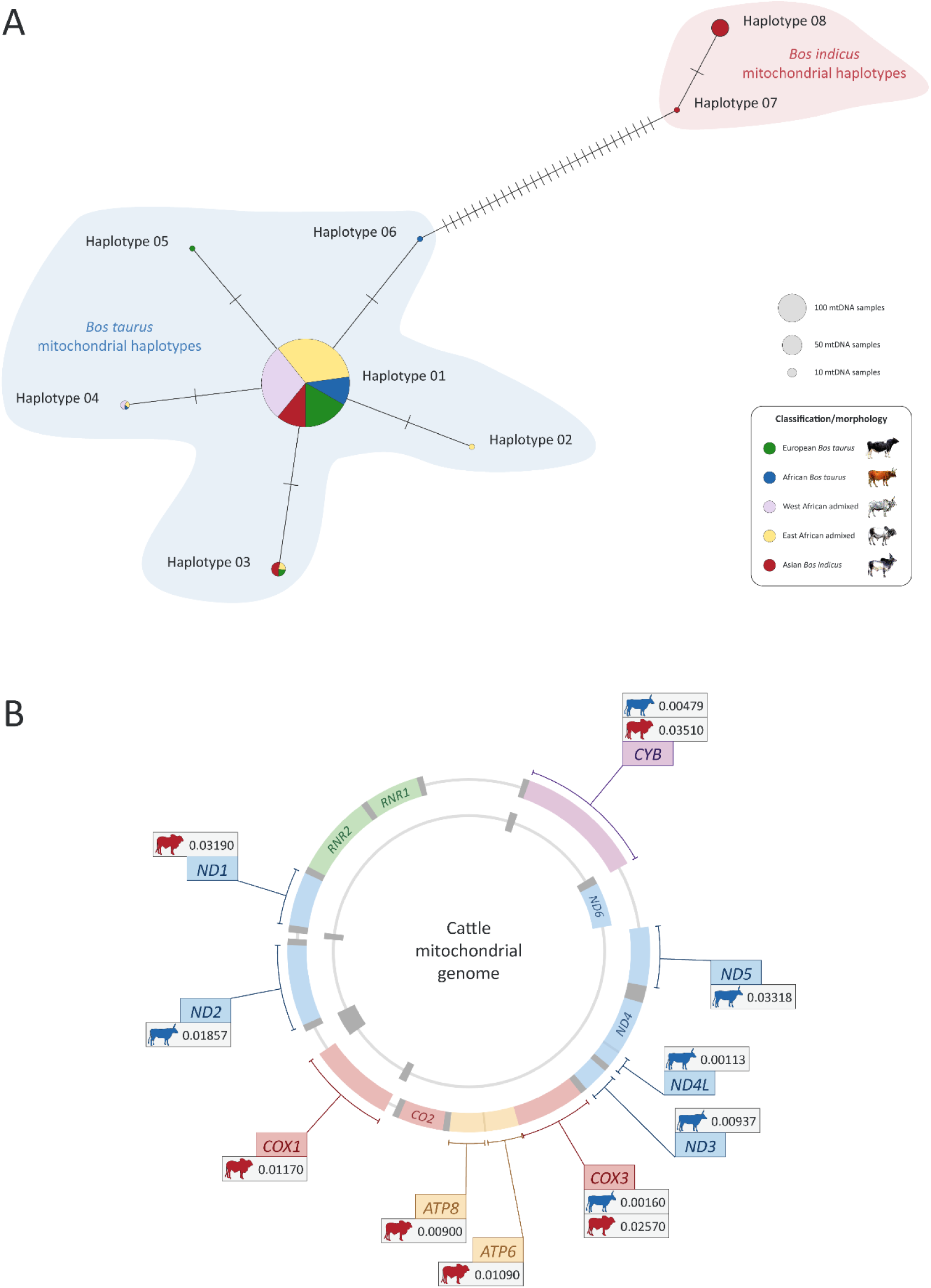
Haplotype diversity and molecular evolution of the cattle mitochondrial genome. (A) Network of 491 cattle mtDNA haplotypes generated using 39 ancestry-informative mtDNA SNPs. This mtDNA haplotype network demonstrates that all surveyed African cattle (47 taurine, 156 East African admixed, and 136 West African admixed) possess Bos taurus mitochondrial genomes. (B) Evidence for positive selection at protein-coding genes in the cattle mitochondrial genome. The significant *P* values (< 0.05) shown in the gene callouts were obtained using the branch-site test of positive selection. The mitochondrially encoded 12S and 16S RNA genes (*RNR1* and *RNR2*) are also shown in green (some figure components created with a <u>BioRender.com</u> license).

### Evidence for positive selection at taurine and zebu mitochondrial OXPHOS protein genes

To assess whether the fixation of taurine mitochondrial ancestry in African cattle could be influenced by mitonuclear incompatibilities, we tested whether bovid mitochondrial sequences possess signals of species-specific adaptation. To do this we obtained high-quality full mtDNA sequences from public DNA sequence databases for 126 African taurine and 21 Asian zebu mitochondrial genomes, and 25 mitochondrial genomes for animals from six additional *Bos* species (*B. gaurus* – gaur; *B. frontalis* – mithun; *B. grunniens* – domestic yak; *B. mutus* – wild yak; *B. javanicus* – banteng; and *B. primigenius* – aurochs) (Table S5). Fixed nucleotide substitutions were identified and catalogued from alignments of the 13 mitochondrial OXPHOS protein gene sequences for African taurine *vs.* Asian zebu, African taurine *vs.* a range of *Bos* species, and Asian zebu *vs.* a range of *Bos* species (Table S2).

We further tested for positive selection at the 13 OXPHOS protein genes using the branch-site test of positive selection (Yang and Nielsen, 2002; Zhang et al., 2005) based on the nonsynonymous/synonymous rate ratio (*ω* = *dN*/*dS*) with positive selection indicated by *ω* > 1 (Table S3). Individual genes showing statistically significant evidence for positive selection are indicated in Figure 3B, which shows that eight of the 13 OXPHOS protein genes have been subject to positive selection in either the taurine (*CYB*, *ND1*, *ND2*, *ND3*, *ND4L*, and *ND5*) or zebu (*ATP6*, *ATP8*, and *COX1*) mitochondrial genomes, and that two (*COX3* and *CYB*) have undergone positive selection in both mtDNA lineages. These results provide strong evidence for positive selection leading to functional differences between zebu and taurine mitochondrial DNA sequences.

### Nuclear-encoded mitochondrially-targeted genes exhibit signatures of coadaptation across admixed African cattle populations

We then assessed whether admixed African cattle populations also preferentially retain taurine ancestry at nuclear genes encoding products targeted to the mitochondrion and those that directly interact with biomolecules produced from the mitochondrial genome. To do this, we reconstructed the local genomic ancestry of East and West African admixed populations, Asian zebu, and African taurine using MOSAIC (Salter-Townshend and Myers, 2019). Three functional subsets of genes were used in this analysis (Table S6): 1) high-confidence “high-mito” genes (HMG) encoding proteins that directly interact with mtDNA-encoded protein subunits in OXPHOS and ribosomal complexes, or that have functions in mtDNA replication (136 genes); 2) lower confidence “low-mito” genes (LMG), which encode proteins that localize to the mitochondrion (661 genes), but are not classified as part of the high-mito subset; and 3) “non-mito” genes (NMG) representing the bulk of the mammalian proteome that does not localize to the mitochondrion (16,383 genes). For each admixed population the taurine and zebu local ancestry estimates were averaged across mitochondrion-targeted genes (the HMG and LMG subsets) and compared to local ancestry estimates from the genomic background (NMG); this produced deviations in taurine local ancestry for each of the three functional gene subsets. We also generated coancestry curve plots using MOSAIC to determine the estimated number of generations since the start of admixture (Figure S2).

From the bootstrap analysis (Figure 4A), we found that three of the ten African admixed breeds individually exhibit significantly more taurine ancestry for the HMG subset: NGAN (*P* = 0.0160), KETE (*P* = 0.0410), EASZ (*P* = 0.0430). Using the non-parametric Wilcoxon signed-rank test across the ten admixed African populations, we also demonstrated that the HMG subset exhibited significant differences in mean taurine ancestries compared to the LMG subset (*P* = 0.0039) and to the NMG subset (*P* = 0.0020). We also compared mean taurine ancestries for the LMG versus the NMG subsets; however, this did produce a significant statistical test result (*P* = 0.2754).

**Figure 4.**
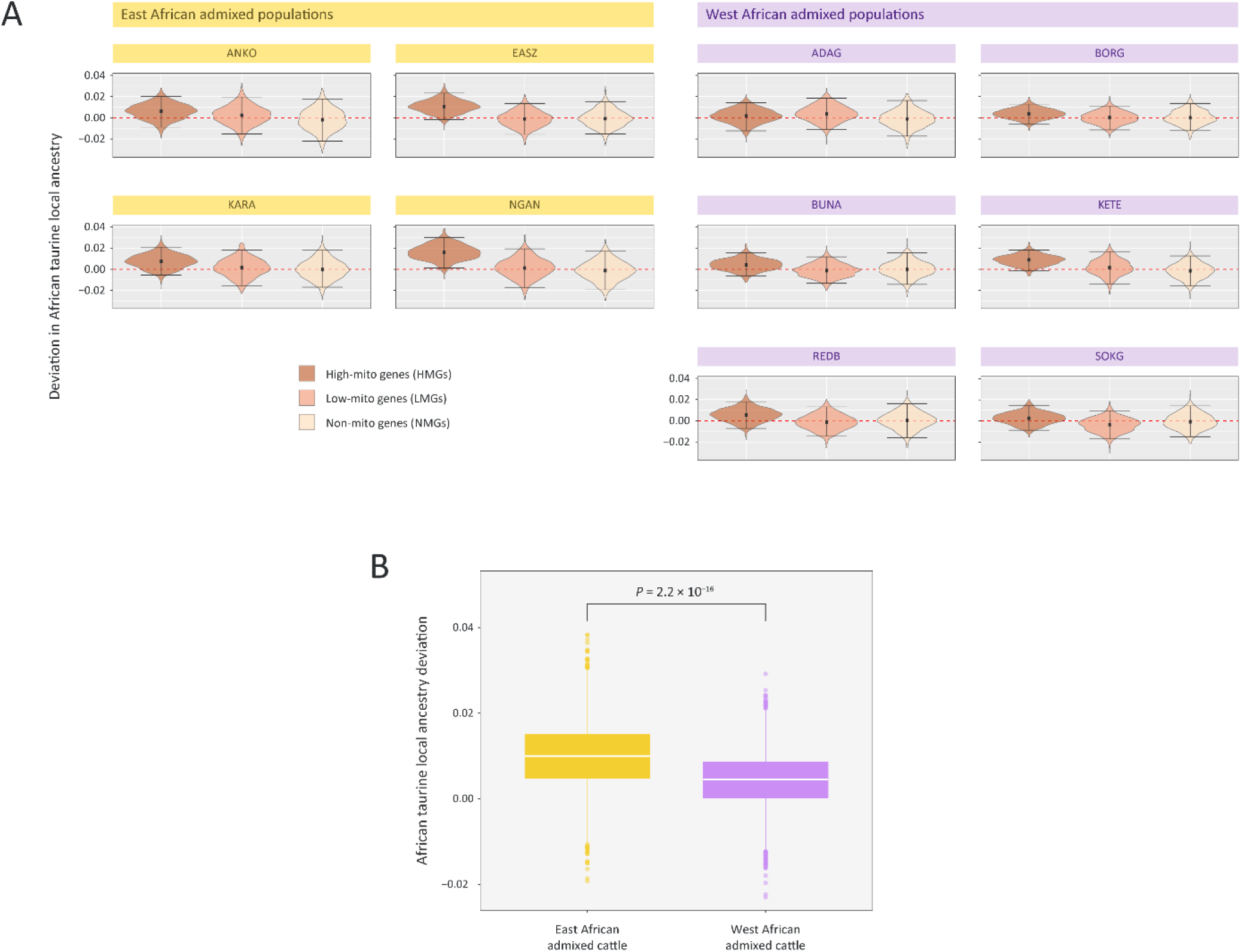
African taurine local ancestry deviations for three different functional gene sets. (A) Violin plots of African taurine local ancestry deviations for the HMG, LMG, and NMG subsets with positive deviations indicating retention of African taurine gene haplotypes. Black data points indicate the median values and horizontal lines represent the 95% confidence interval. (*B*) Box plot of African taurine local ancestry deviations for the HMG subset in the East African and the West African admixed groups. White lines indicate the median values and yellow and purple boxes indicate the interquartile ranges.

### Functional consequences of mitonuclear incompatibilities in admixed African cattle breeds

Previous studies have examined sub-chromosomal admixture and local ancestry in hybrid taurine/zebu animals (Barbato et al., 2020; Chen *et al*., 2018; Koufariotis et al., 2018; McTavish and Hillis, 2014), and we extend this work to mitonuclear incompatibilities and coadaptation in admixed cattle populations. Using a high-density SNP genotyping array, ten different breeds were examined with genome-wide zebu ancestries ranging between 37% (Borgou) and 74% (Karamojong), and estimated dates for the start of admixture in each population extending from the 14^th^ to the 20^th^ century (Figure S1 and Table S1). A consistent pattern of mitonuclear disequilibria was observed for the functional HMG subset within three breeds of admixed African cattle (EASZ, KETE, NGAN) (Figure 4A): African taurine local ancestry was uniformly higher for nuclear genes encoding proteins that directly engage with mitochondrial-encoded gene products to form multi-subunit complexes, or that directly interact with mitochondrial DNA or RNAs; this subset encompasses genes that encode OXPHOS subunits, ribosomal proteins, tRNA synthetases, and DNA and RNA polymerases. In support of the hypothesis that functional incompatibilities exist between the taurine and zebu mitochondrial genomes, we also find compelling evidence that the two mtDNA lineages have been subject to positive selection at ten of the 13 OXPHOS protein genes (Figure 3B and Table S2).

Although the source population divergence is substantially less in admixed humans, these results are comparable to those obtained by Zaidi and Makova (2019), which support the hypothesis that selection in admixed human populations has acted against mitonuclear incompatibilities. They observed significant enrichment of sub-Saharan African ancestry for HMG subset genes in an African American population with sub-Saharan African and European nuclear ancestry and predominantly sub-Saharan African mtDNA haplotypes. They also observed significant enrichment of Native American ancestry at HMG subset genes in a Puerto Rican population with Native American and European nuclear ancestry and predominantly Native American mtDNA haplotypes.

The functional HMG and LMG subsets containing 136 and 661 genes, respectively (Table S6) were used in the present study for the purpose of evaluating mitonuclear incompatibilities in admixed African cattle populations. However, it is also instructive to examine these genes in the context of recently published high-resolution surveys of African cattle genomic diversity and signatures of selection (Table S4). For example, the aspartyl-tRNA synthetase 2, mitochondrial gene (*DARS2*), an HMG subset gene on BTA16, is in the region encompassed by selective sweeps detected separately in the EASZ breed and a composite sample of East African zebu cattle (Bahbahani et al., 2017; Taye et al., 2018). Inspection of the Cattle Gene Atlas (Fang et al., 2020) demonstrates that *DARS2* is most highly expressed in spermatozoa and therefore functionally linked to sperm motility, which may provide an explanation for mitonuclear coevolution in admixed cattle at this locus. In addition, the mitochondrial ribosomal protein S33 gene (*MRPS33*), another HMG subset gene, was detected within a positively selected region on BTA4 when African cattle were compared to commercial European and Asian breeds (Kim *et al*., 2017) and in analyses of selective sweeps focused on the evolution of thermotolerance in African cattle populations (Taye et al., 2017).

Agriculture in Sub-Saharan Africa rely on a diverse array of indigenous cattle breeds, many of which show marked resilience to harsh environments, climatic extremes, and infectious disease—adaptations that have been shaped by their dual taurine-zebu ancestry. Cattle breeding programs in Africa are currently poised to leverage this composite ancestry through genomic selection as a leapfrog technology to bypass conventional breeding to enhance resilience (e.g., via the superior thermotolerance of zebu cattle), production, health, and welfare traits and ultimately improve the livelihoods of smallholder farmers (Ibeagha-Awemu et al., 2019; Marshall et al., 2019; Mrode et al., 2019). Future genetic improvement programs in African cattle will therefore need to consider mitonuclear incompatibilities that could reduce the fitness of hybrid taurine/zebu breeds. Understanding these incompatibilities in hybrid cattle may also provide useful information for targeted editing of both the bovine mitochondrial genome and mitochondrially-targeted genes in the nuclear genome (Klucnika and Ma, 2020; Tang et al., 2021). Finally, our results demonstrate that admixed African cattle populations can serve as comparative model systems for understanding the phenotypic consequences of mitonuclear interactions and adaptive and maladaptive genomic introgression in other mammals, including humans.

### Limitations of the study

Although we provide support for the hypothesis that mitonuclear coevolution exists between the nuclear and mitochondrial genomes of hybrid African cattle populations, this work is necessarily limited by the number of populations sampled and the density of the SNP data used. In addition, the genome-wide approach we use here is not directly amenable to gene-by-gene analyses, which could use whole-genome sequence data sets from large numbers of hybrid animals to directly identify incompatibilities between individual nuclear- and mitochondrial-encoded proteins.

## SUPPLEMENTAL INFORMATION

Supplemental information can be found online at https://doi.org [to be determined].

## ACKNOWLEDGEMENTS

J.A.W was supported by Science Foundation Ireland (SFI) and Acceligen/Recombinetics Inc. through the SFI Centre for Research Training in Genomics Data Science under grant no. 18/CRT/6214. G.P.M., T.J.H., and D.E.M. were supported by an SFI Investigator Award (SFI/15/IA/3154). T.S.S. and D.E.M were supported by a United States Department of Agriculture (USDA) and Department of Agriculture, Food, and the Marine (DAFM) US-Ireland Research and Development Partnership grant (TARGET-TB, 17/RD/US-ROI/52).

## AUTHOR CONTRIBUTIONS

Conceptualization, J.A.W., M.J.D., T.S.S., D.G.B., L.A.F., M.S-T., and D.E.M.; investigation, J.A.W., G.P.M., T.J.H, S.I.N, L.A.F., M.S-T., and D.E.M.; formal analysis: J.A.W., G.P.M., T.J.H, and S.I.I.N; data curation, J.A.W., S.I.N, L.A.F., and D.E.M.; resources, T.S.S., D.G.B., L.A.F., and D.E.M.; writing – original draft, J.A.W. and D.E.M.; writing – review and editing, all authors; visualization, J.A.W., S.I.N, and D.E.M.; supervision: L.A.F., M.S-T., and D.E.M.; funding acquisition, T.S.S., L.A.F., M.S-T., and D.E.M.

## STAR★METHODS

### RESOURCE AVAILABILITY

#### Lead contact

Further information and inquiries about code, reagents and/or data details may be directed to the lead contact, David E. MacHugh.

#### Materials availability

This study did not generate new unique reagents.

#### Data and code availability

This study did not generate any unpublished custom code, software, or algorithm. New Illumina^®^ BovineHD 777K BeadChip SNP data generated for this study is available from the Dryad Digital Repository (https://datadryad.org) [DOI accession link to be provided prior to publication].

### EXPERIMENTAL MODEL, AND SUBJECT DETAILS

#### Animal sampling and genotyping

High-density genome-wide SNP array data sets (Illumina^®^ BovineHD 777K BeadChip) corresponding to a total of 605 animals were obtained from published studies (Bahbahani et al., 2015; Verdugo *et al*., 2019) and collated in the Acceligen cattle genotyping database. For the present study, new BovineHD 777K BeadChip SNP data were generated for 73 additional animals (50 Borgou and 23 N’Dama) that were previously published by our group as part of microsatellite-based surveys of cattle genetic diversity (Freeman et al., 2004; MacHugh *et al*., 1997). These new SNP genotype data sets were generated by Weatherbys Scientific (Co. Kildare, Ireland) using standard procedures for Illumina SNP array genotyping. In total, 18 different breeds/populations were represented (Table 1), including two West African taurine breeds (Muturu and N’Dama); two European taurine breeds (Holstein-Friesian and Jersey); ten West and East African admixed zebu-taurine (Adamawa Gudali, Ankole, Borgou, Bunaji, East African Shorthorn Zebu, Karamojong, Keteku, Nganda, Red Bororo, and Sokoto Gudali); and four zebu breeds of South Asian origin (Gir, Hariana, Nelore, and Sahiwal). Table 1 also shows the three- or four-letter codes used to designate each breed.

#### SNP data formatting and quality control

BovineHD 777K SNP locations were remapped to the current bovine genome assembly ARS-UCD1.2 (Rosen et al., 2020) and SNP genotype data were merged using PLINK v1.9 (Chang et al., 2015). Quality control (QC) of the combined SNP genotype data set was also performed using PLINK v1.9 and autosomal SNPs with a call rate < 95% and a minor allele frequency (MAF) of < 0.05 were filtered from the data.

### QUANTIFICATION AND STATISTICAL ANALYSIS

#### Principal component and structure analyses

Principal component analysis (PCA) of individual animal SNP genotype data for the African taurine (MUTU and NDAG), the East African admixed (ADAG, ANKO, BORG, BUNA, EASZ, KARA, KETE, NGAN, REDB, and SOKG) and two Asian indicine (GIR and NELO) populations was performed using PLINK v1.9 and the results were plotted using ggplot2 v3.3.3 (Wickham, 2016) in the R v3.6.2 environment for statistical computation and graphics (R Core Team, 2019). The genetic structure of each population was also estimated using fastSTRUCTURE v1.0 with *K* = 3 modelled ancestries to determine mean African taurine, European taurine, and Asian zebu contributions (Raj et al., 2014).

#### Mitochondrial DNA haplogroup determination

The BovineHD 777K BeadChip includes 346 SNPs located in the mitochondrial genome, which can be used to construct haplotypes and catalogue and distinguish the mitochondrial haplogroups characteristic of *B. taurus* and *B. indicus* cattle lineages. For this analysis, the European JRSY and HOLS taurine breeds and the Indo-Pakistan HARI and SAHI Asian indicine breeds were also included to ensure good representation of the *B. taurus* and *B. indicus* mtDNA haplogroups—the ‘T’ and ‘I’ groups, respectively (Chen et al., 2010; Troy *et al*., 2001). The mtDNA SNPs were filtered using PLINK v1.9 (Chang *et al*., 2015) such that SNPs with a MAF of < 0.10, and a call rate of < 95% were removed. Individual animals with a genotype missingness of > 95% were also removed. Following this, the most ancestry informative mtDNA SNPs were identified using infocalc (Rosenberg, 2005; Rosenberg et al., 2003), which provides *I*_n_, a general measure of the informativeness of a SNP for ancestry assignment. The 50 top ranked SNPs, based on *I*_n_, were then used to generate mtDNA haplotypes with the fastPHASE v1.4 program (Scheet and Stephens, 2006). Haplotype networks were constructed using the POPART v1.7 package (Leigh and Bryant, 2015).

#### Molecular evolution of mtDNA OXPHOS genes

Complete mitochondrial genome sequences for three groups of cattle and related species were obtained from publicly available DNA sequence databases (Table S5). The mitochondrial genome sequences used represented the African *B. taurus* (126 animals), and Asian *B. indicus* (21 animals) mtDNA lineages, and the following additional *Bos* species: *B. gaurus* – gaur (6 animals); *B. frontalis* – mithun (4 animals); *B. grunniens* – domestic yak (5 animals); *B. mutus* – wild yak (4 animals); *B. javanicus* – banteng (4 animals); *and B. primigenius* – aurochs (2 animals). The protein-coding sequence for 13 essential OXPHOS genes were aligned using the MAFFT v.7.49 software package (Katoh et al., 2019). Evidence for positive selection at the 13 OXPHO protein genes (*ATP6*, *ATP8*, *CYB*, *COX1*, *COX2*, *COX3*, *ND1*, *ND2*, *ND3*, *ND4*, *ND4L*, *ND5*, and *ND6*) was evaluated using the *d*_N_/*d*_S_ ratio (*ω*) branch site test for positive selection (Yang and Nielsen, 2002; Zhang *et al*., 2005) with the CODEML branch-site models MA( ω > 1) *vs.* MA( ω = 1) implemented in the PAML v4.9 software package (Yang, 2007).

#### Local ancestry analysis of admixed populations

Local ancestry across the bovine genome for each African admixed breed (ADAG, ANKO, BORG, BUNA, EASZ, KARA, KETE, NGAN, REDB, and SOKG) was inferred using MOSAIC v1.3.7 (Salter-Townshend and Myers, 2019). The MOSAIC algorithm, unlike other methods, does not require defined surrogate donor reference populations for the mixing ancestral populations; it fits a two-layer Hidden Markov Model (HMM) that determines how closely related each segment of chromosome in each admixed individual genome is to the segments of chromosomes in individual genomes from potential donor populations. While determining local ancestry along each chromosome, MOSAIC also infers the number of generations since the admixture process started for a particular population. The potential donor populations used for the MOSAIC local ancestry analysis were the two West African *B. taurus* breeds (MUTU and NDAG) and two of the Asian *B. indicus* breeds (GIR and NELO). The MOSAIC algorithm requires phased haplotypes and a recombination rate map; therefore, SHAPEIT v2 (r900) (Delaneau et al., 2012) was used to generate phased haplotypes and a published cattle recombination map was employed (Martin et al., 2015).

#### Detection of taurine local ancestry deviation

To determine if there was significant retention of *B*. *taurus* nuclear genes that encode mitochondrially targeted proteins (*N-mito* genes) in African admixed cattle, we used an approach modified from previously published surveys of mitonuclear incompatibilities in modern admixed human populations (Sloan et al., 2015; Zaidi and Makova, 2019) and because of ancient gene flow from archaic hominins (*H. neanderthalensis* and *H. denisova*) (Sharbrough *et al*., 2017). Firstly, the MitoCarta 2.0 database resource (Calvo et al., 2016) was used to obtain an inventory of genes that produce the nuclear-encoded component of the mammalian mitochondrial proteome, i.e., proteins with experimental evidence for localization in the mitochondrion. Following this, the Ensembl BioMart tool (Yates et al., 2020) was used to generate a list of 1158 bovine N-mito genes, which was classified into two functional subsets as defined by Sloan *et al*. (2015) and also used by Sharbrough *et al*. (2017) and Zaidi and Makova (2019). These subsets were denoted as 1) high-confidence “*high-mito*” genes (HMG) encoding proteins that directly interact with mtDNA-encoded protein subunits in OXPHOS and ribosomal complexes, or that have functions in mtDNA replication (136 genes); and 2) lower confidence “*low-mito*” genes (LMG), which encode proteins that localize to the mitochondrion (661 genes) but are not classified as part of the high-mito subset. Finally, a third group of “*non-mito*” genes (NMG) was generated, which includes the bulk of the mammalian proteome that does not localize to the mitochondrion (16,383 genes). Table S6 provides further detail for the functional gene subsets used to detect evidence for mitonuclear incompatibilities in African admixed cattle populations.

The local ancestry estimates generated using MOSAIC for each SNP across the genome were catalogued and the BEDTools v2.18 software suite (Quinlan and Hall, 2010) was then used to intersect these SNPs with windows spanning 2.5 Mb upstream and downstream of genes within each of the three functional gene subsets. Following this, and as described by Zaidi and Makova (2019), for each of the three subsets an unweighted block bootstrap approach was used to generate this methodology is subtraction of the mean ancestry fraction across the local ancestry estimate for each SNP (the *expectation*), which produces the deviation in local ancestry at each SNP locus. For each functional gene subset, the number of windows sampled with replacement was the same as the number of HMG subset genes (*n* = 136). In each case, the mean ancestry deviation was estimated and then averaged across all windows. Bootstrap resampling (1000 replicates) was used to generate a distribution of mean deviations in local ancestry for each of the three functional gene subsets. Overall significance of the distributions was assessed by the proportion of the distribution that overlapped zero. Mean taurine ancestry was determined for each of the gene subsets across all ten populations. Differences in the population means of these were assessed using the non-parametric Wilcoxon signed-rank test in R v3.6.2 (R Core Team, 2019).

## KEY RESOURCES TABLE

The table highlights the reagents, genetically modified organisms and strains, cell lines, software, instrumentation, and source data **essential** to reproduce results presented in the manuscript. Depending on the nature of the study, this may include standard laboratory materials (i.e., food chow for metabolism studies, support material for catalysis studies), but the table is **not** meant to be a comprehensive list of all materials and resources used (e.g., essential chemicals such as standard solvents, SDS, sucrose, or standard culture media do not need to be listed in the table). **Items in the table must also be reported in the method details section within the context of their use.** To maximize readability, the number of **oligonucleotides and RNA sequences** that may be listed in the table is restricted to no more than 10 each. If there are more than 10 oligonucleotides or RNA sequences to report, please provide this information as a supplementary document and reference the file (e.g., See Table S1 for XX) in the key resources table.

***Please note that ALL references cited in the key resources table must be included in the references list***. Please report the information as follows:

• **REAGENT or RESOURCE:** Provide full descriptive name of the item so that it can be identified and linked with its description in the manuscript (e.g., provide version number for software, host source for antibody, strain name). In the experimental models section (applicable only to experimental life science studies), please include all models used in the paper and describe each line/strain as: model organism: name used for strain/line in paper: genotype. (i.e., Mouse: OXTR^fl/fl^: B6.129(SJL)- Oxtr^tm1.1Wsy/J^). In the biological samples section (applicable only to experimental life science studies), please list all samples obtained from commercial sources or biological repositories. Please note that software mentioned in the methods details or data and code availability section needs to also be included in the table. See the sample tables at the end of this document for examples of how to report reagents.
• **SOURCE:** Report the company, manufacturer, or individual that provided the item or where the item can be obtained (e.g., stock center or repository). For materials distributed by Addgene, please cite the article describing the plasmid and include “Addgene” as part of the identifier. If an item is from another lab, please include the name of the principal investigator and a citation if it has been previously published. If the material is being reported for the first time in the current paper, please indicate as “this paper.” For software, please provide the company name if it is commercially available or cite the paper in which it has been initially described.
• **IDENTIFIER:** Include catalog numbers (entered in the column as “Cat#” followed by the number, e.g., Cat#3879S). Where available, please include unique entities such as <u>RRIDs</u>, Model Organism Database numbers, accession numbers, and PDB, CAS, or CCDC IDs. For antibodies, if applicable and available, please also include the lot number or clone identity. For software or data resources, please include the URL where the resource can be downloaded. Please ensure accuracy of the identifiers, as they are essential for generation of hyperlinks to external sources when available. Please see the Elsevier <u>list of data repositories</u> with automated bidirectional linking for details. When listing more than one identifier for the same item, use semicolons to separate them (e.g., Cat#3879S; RRID: AB_2255011). If an identifier is not available, please enter “N/A” in the column.

`o` ***A NOTE ABOUT RRIDs:*** We highly recommend using RRIDs as the identifier (in particular for antibodies and organisms but also for software tools and databases). For more details on how to obtain or generate an RRID for existing or newly generated resources, please <u>visit the RII</u> or <u>search for RRIDs</u>.

Please use the empty table that follows to organize the information in the sections defined by the subheading, skipping sections not relevant to your study. Please do not add subheadings. To add a row, place the cursor at the end of the row above where you would like to add the row, just outside the right border of the table. Then press the ENTER key to add the row. Please delete empty rows. Each entry must be on a separate row; do not list multiple items in a single table cell. Please see the sample tables at the end of this document for relevant examples in the life and physical sciences of how reagents and instrumentation should be cited.

## TABLE FOR AUTHOR TO COMPLETE

Please upload the completed table as a separate document. **<u>Please do not add subheadings to the key resources</u> <u>table.</u>** If you wish to make an entry that does not fall into one of the subheadings below, please contact your handling editor. **<u>Any subheadings not relevant to your study can be skipped.</u>** (**NOTE:** For authors publishing in Cell Genomics, Cell Reports Medicine, Current Biology, and Med, please note that references within the KRT should be in numbered style rather than Harvard.)

### Key resources table

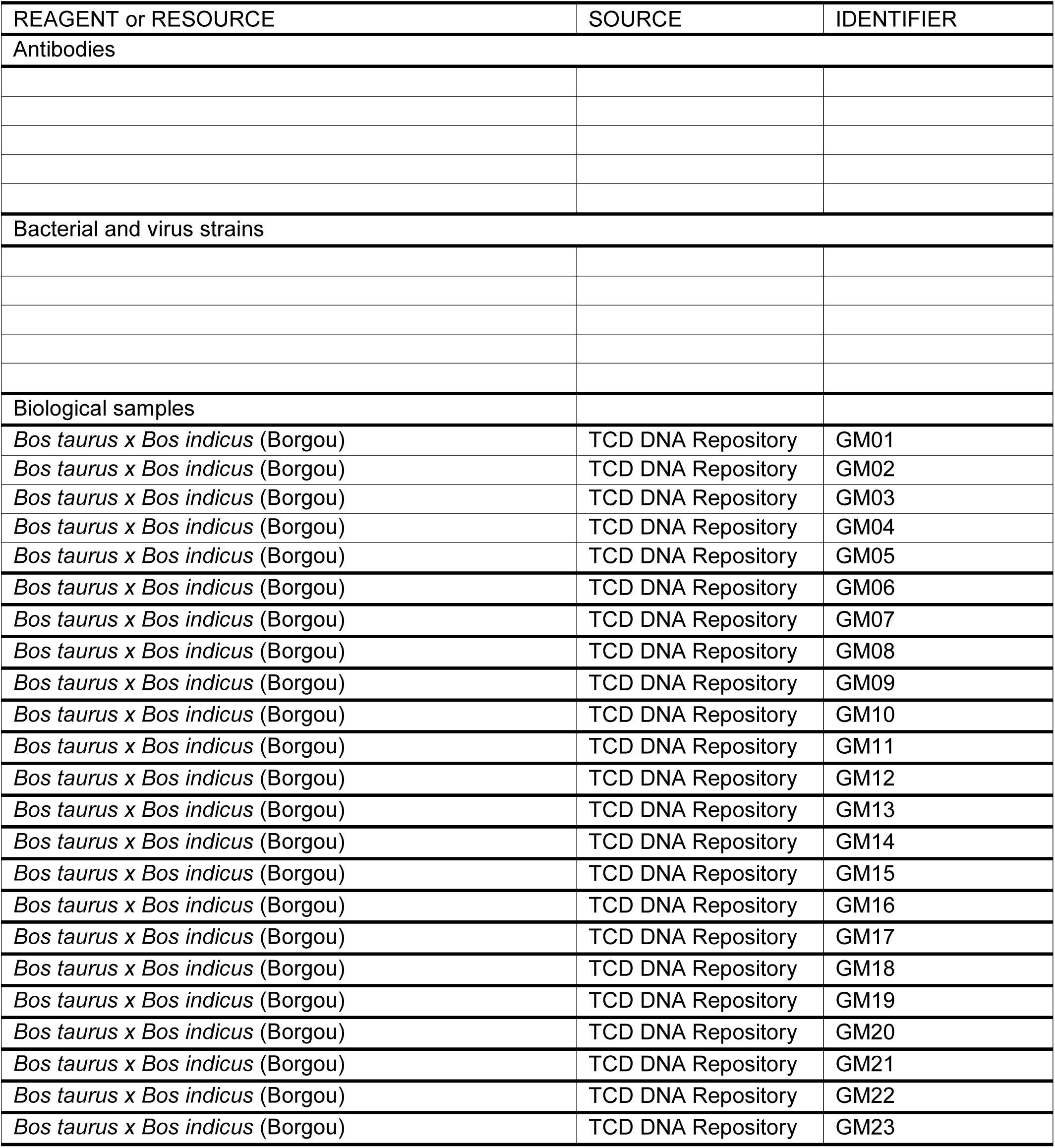
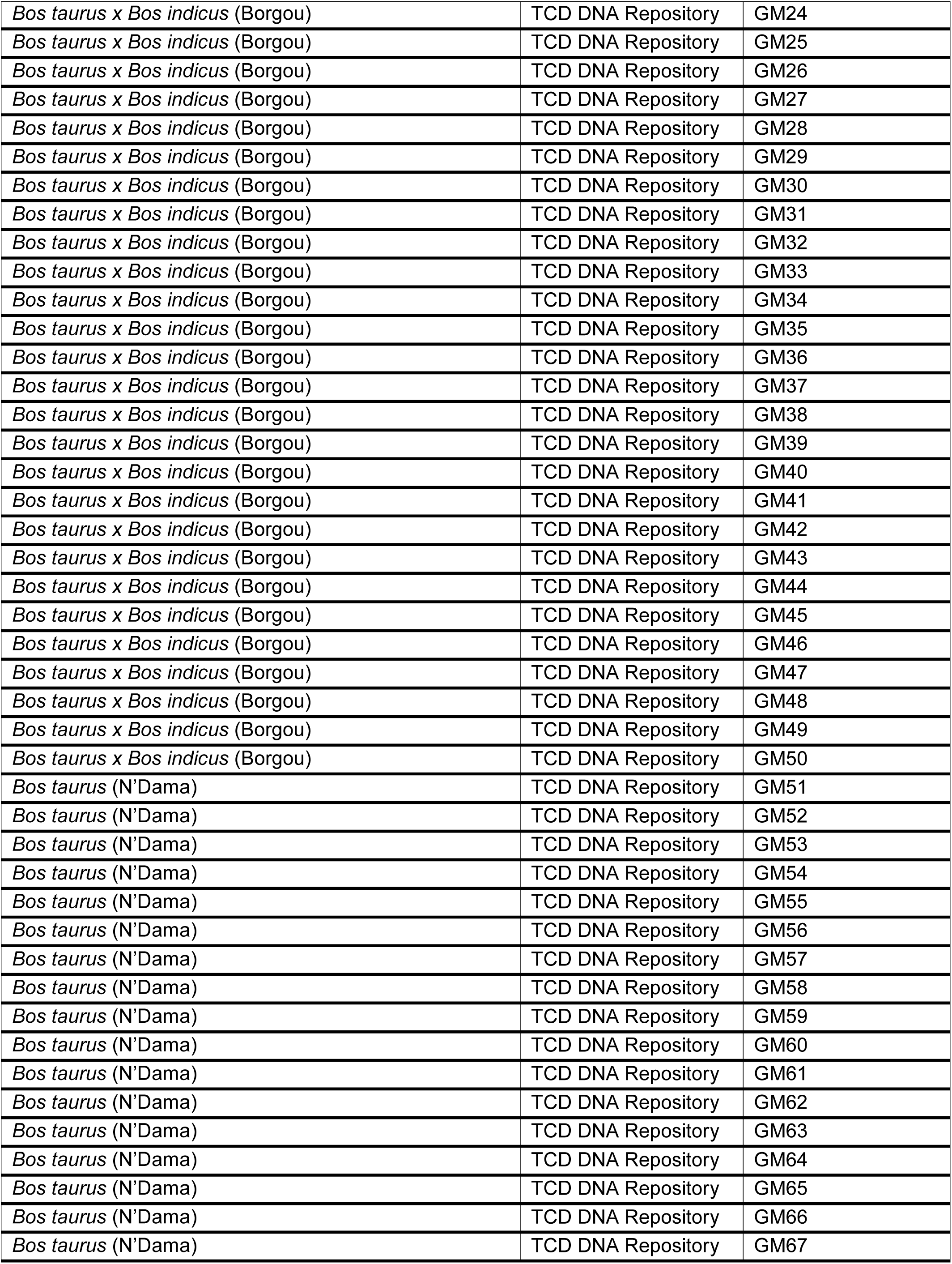
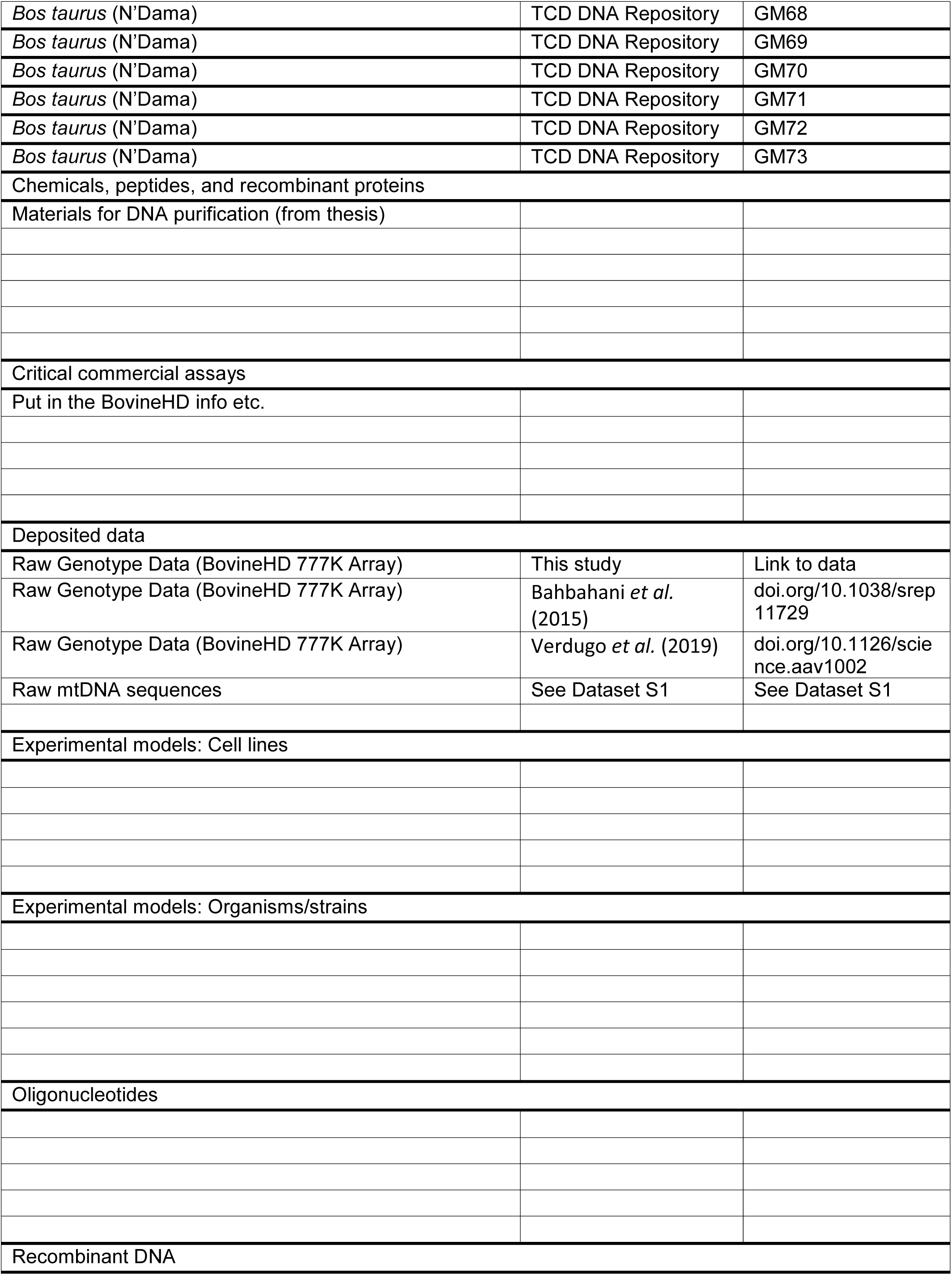
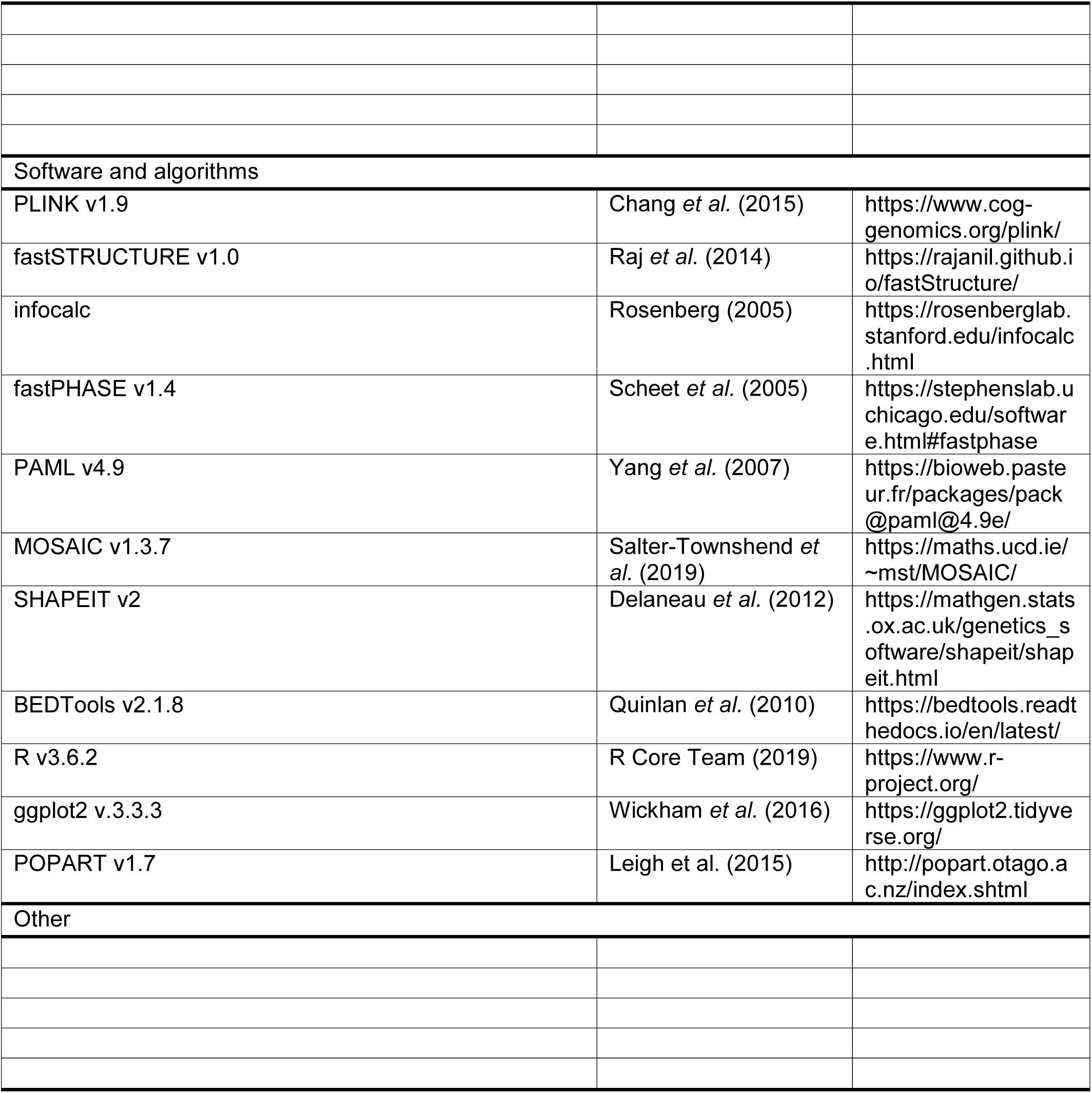

## Supplementary Information

**Figure S1.**
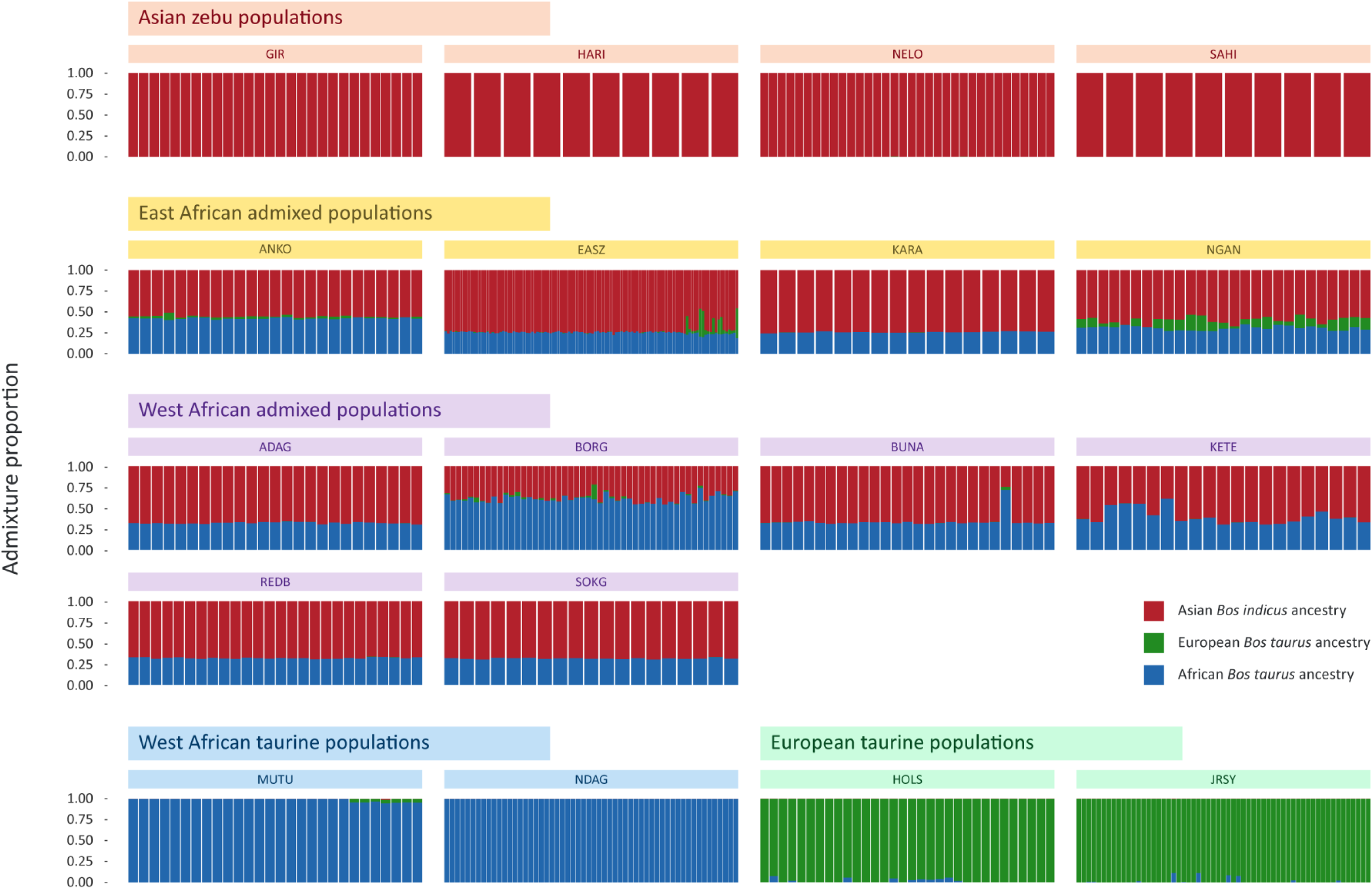
Unsupervised genetic structure plot for Asian zebu, East and West African admixed cattle, and West African and European taurine breeds. Results for an inferred number of ancestry clusters of *K* = 3 is shown, which corresponds to Asian *Bos indicus* (red), European *Bos taurus* (green), and African *B. taurus* (blue) ancestral components, respectively.

**Figure S2.**
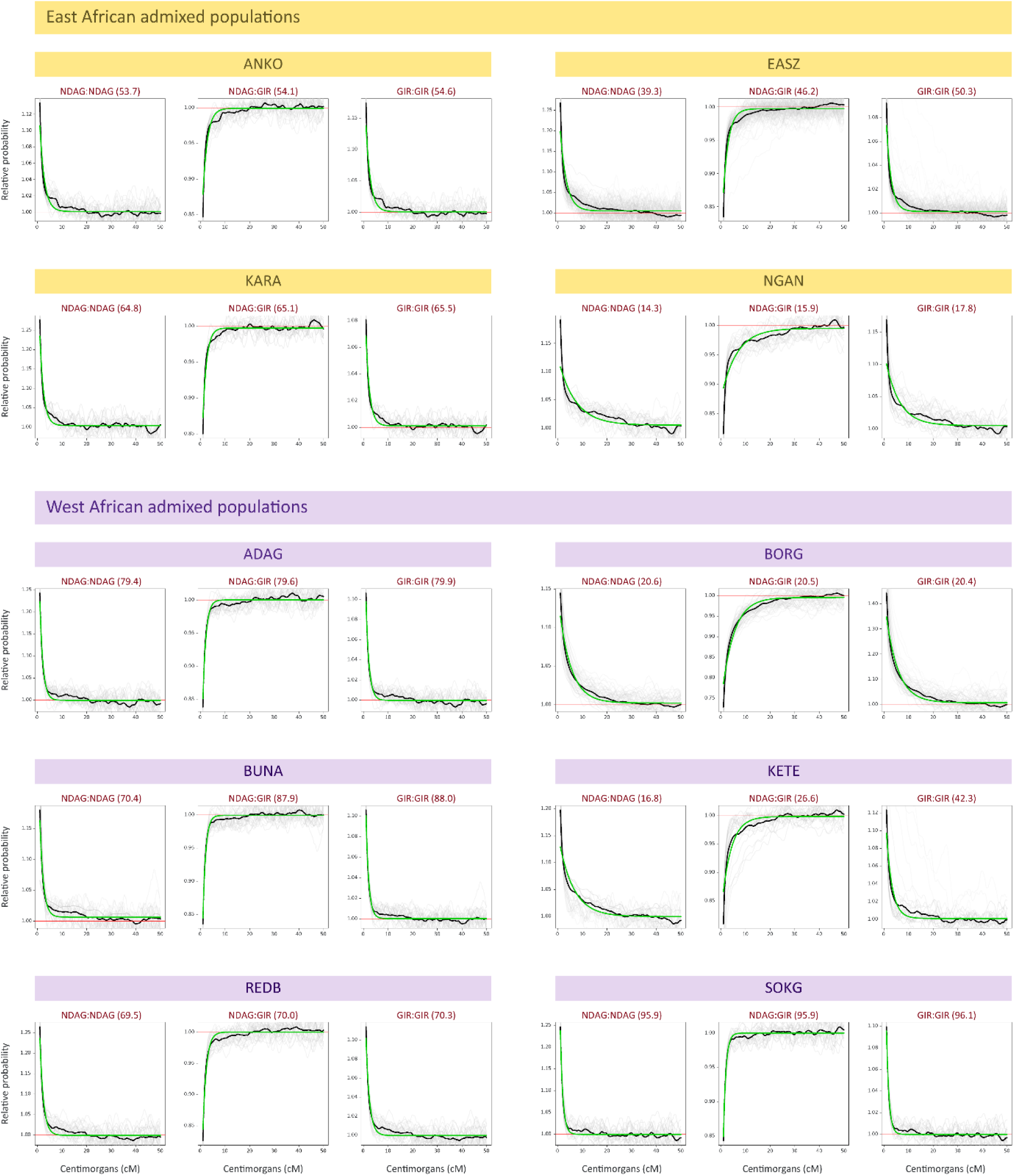
Coancestry curve plots generated using MOSAIC for 10 East and West African admixed cattle populations. These curves show the exponential decay of the ratio of probabilities of pairs of local ancestries (y-axis) as a function of genetic distance (x-axis). The pair of ancestries used for each curve is shown on the top of each plot with the estimated number of generations since the start of admixture in brackets. For each plot, the green line represents the fitted curve, the black line shows the across targets ratio, and the grey lines indicate the per target ratio (further information in Salter-Townshend and Myers, 2019).

**Table S1.** Ancestry components estimated using fastSTRUCTURE with estimated times for the start of the admixture process in African admixed cattle populations generated using MOSAIC.

| Code | Type/morphology | Country of origin | African taurine ancestry | European taurine ancestry | Asian zebu ancestry | Generations since start of admixture | Start of admixture |
| --- | --- | --- | --- | --- | --- | --- | --- |
| MUTU | West African taurine | Nigeria | 0.990 ± 0.019 | 0.010 ± 0.018 | 0.000 ± 0.003 |  | ---- |
| NDAG | West African taurine | Guinea | 1.000 ± 0.000 | 0.000 ± 0.000 | 0.000 ± 0.000 |  | ---- |
| HOLS | European taurine | Netherlands | 0.010 ± 0.025 | 0.990 ± 0.025 | 0.000 ± 0.000 |  | ---- |
| JRSY | European taurine | United Kingdom | 0.010 ± 0.022 | 0.990 ± 0.022 | 0.000 ± 0.000 |  | ---- |
| ANKO | East African admixed | Uganda | 0.420 ± 0.010 | 0.020 ± 0.016 | 0.550 ± 0.013 | 54.1 | 1641 – 1804 CE |
| EASZ | East African admixed | Kenya | 0.250 ± 0.015 | 0.020 ± 0.065 | 0.730 ± 0.055 | 46.2 | 1697 – 1835 CE |
| KARA | East African admixed | Uganda | 0.260 ± 0.008 | 0.000 ± 0.001 | 0.740 ± 0.008 | 65.1 | 1564 – 1760 CE |
| NGAN | East African admixed | Uganda | 0.310 ± 0.024 | 0.100 ± 0.052 | 0.590 ± 0.039 | 15.9 | 1909 – 1956 CE |
| ADAG | West African admixed | Nigeria | 0.320 ± 0.010 | 0.000 ± 0.001 | 0.680 ± 0.010 | 79.6 | 1463 – 1702 CE |
| BORG | West African admixed | Benin | 0.610 ± 0.047 | 0.010 ± 0.027 | 0.370 ± 0.055 | 20.5 | 1877 – 1938 CE |
| BUNA | West African admixed | Nigeria | $0.340 \pm 0.076$ | $0.000 \pm 0.007$ | $0.660 \pm 0.083$ | 87.9 | 1405 – 1668 CE |
| KETE | West African admixed | Nigeria | $0.400 \pm 0.093$ | $0.000 \pm 0.001$ | $0.600 \pm 0.093$ | 26.6 | 1834 – 1914 CE |
| REDB | West African admixed | Nigeria | $0.320 \pm 0.009$ | $0.000 \pm 0.001$ | $0.680 \pm 0.009$ | 70 | 1530 – 1740 CE |
| SOKG | West African admixed | Nigeria | $0.320 \pm 0.008$ | $0.000 \pm 0.000$ | $0.680 \pm 0.008$ | 95.9 | 1349 – 1636 CE |
| GIR | Asian zebu | India | $0.000 \pm 0.000$ | $0.000 \pm 0.000$ | $1.000 \pm 0.000$ | | ---- |
| HARI | Asian zebu | India | $0.000 \pm 0.000$ | $0.000 \pm 0.000$ | $1.000 \pm 0.000$ | | ---- |
| NELO | Asian zebu | Brazil | $0.000 \pm 0.000$ | $0.000 \pm 0.001$ | $1.000 \pm 0.001$ | | ---- |
| SAHI | Asian zebu | India | $0.000 \pm 0.000$ | $0.000 \pm 0.000$ | $1.000 \pm 0.000$ | | ---- |

*Note.* For each admixed population, the generations since admixture started were obtained from the NDAG:GIR coancestry plots (Fig. S2). A generation interval range of 4–7 years for managed domestic cattle was used. CE = Common Era.

**Table S2.**
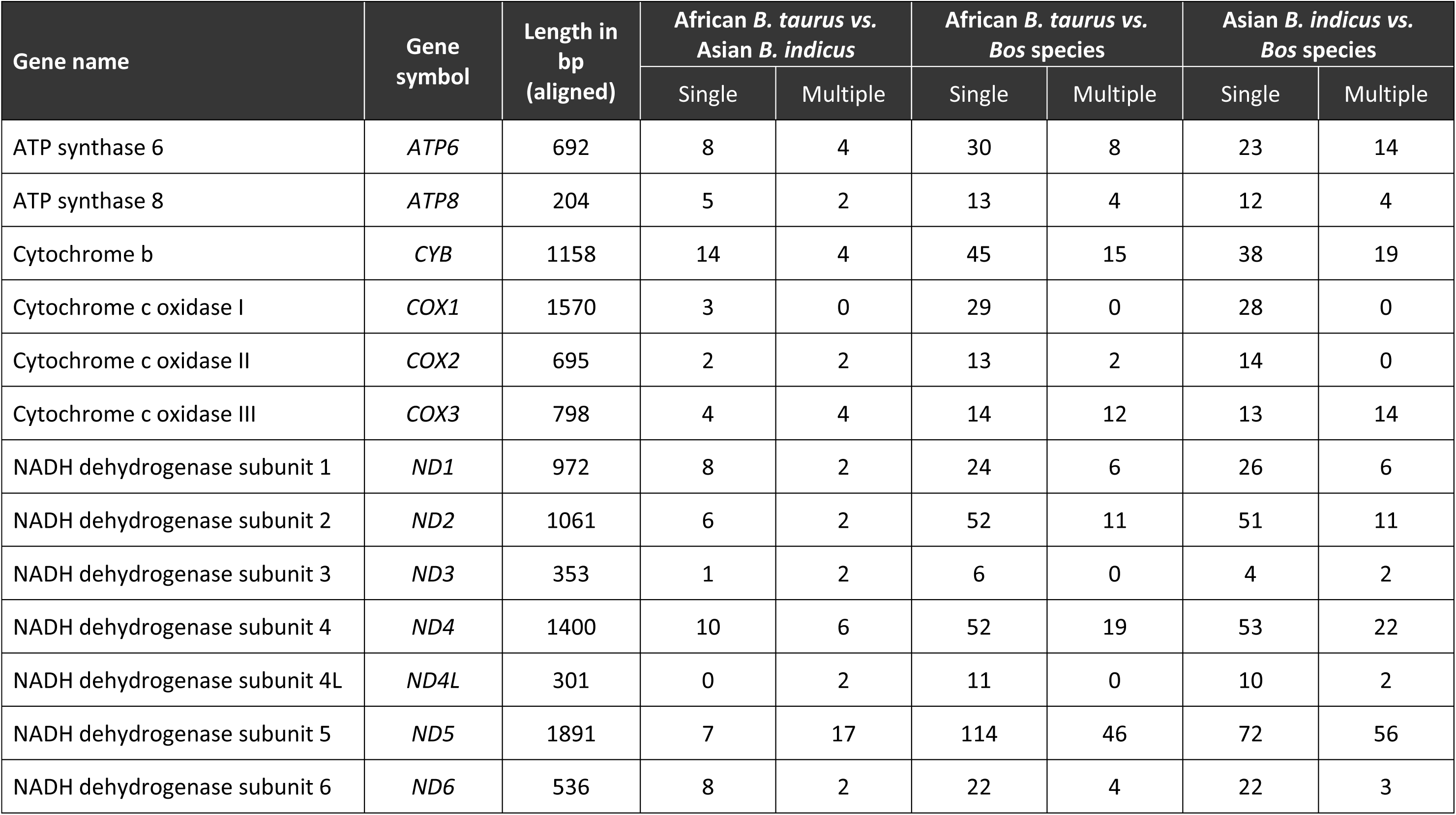
Fixed nucleotide substitutions determined from alignments of the protein-coding sequences of 13 mitochondrial OXPHOS protein genes for three groups of Bovinae species/subspecies. The taxa examined included African *Bos taurus*, Asian *Bos indicus* and a range of *Bos* species (*Bos gaurus*, *Bos grunniens*, *Bos javanicus*, *Bos mutus, Bos frontalis, and Bos primigenius*).

**Table S3.** Results for the branch-site test of positive selection (BSPS) for 13 mitochondrial OXPHOS protein gene sequences in three different *Bos* groups. Significant *P* values (< 0.05) indicating positive selection for individual genes are shown in bold underline.

| *Bos taurus* mitochondrial genome sequences |  |  |  |  |  |  |  |
| --- | --- | --- | --- | --- | --- | --- | --- |
| Gene symbol | No. param. $H_A$ | No. param. $H_0$ | df | LnL $H_A$ | LnL $H_0$ | LRT(2ΔLnL) | *P* ($\chi^2$) |
| *ATP6* | 381 | 380 | 1 | -2101.88 | -2101.79 | -0.18280 | 1.00000 |
| *ATP8* | 381 | 380 | 1 | -527.49 | -526.98 | -1.01728 | 1.00000 |
| *CYB* | 381 | 380 | 1 | -3334.59 | -3334.59 | 0.00004 | **<u>0.00479</u>** |
| *COX1* | 381 | 380 | 1 | -3955.67 | -3955.67 | -0.00081 | 1.00000 |
| *COX2* | 381 | 380 | 1 | -1696.07 | -1696.06 | -0.01870 | 1.00000 |
| *COX3* | 381 | 380 | 1 | -1774.40 | -1774.40 | 0.00000 | **<u>0.00160</u>** |
| *ND1* | 381 | 380 | 1 | -2475.83 | -2475.83 | -0.00070 | 1.00000 |
| *ND2* | 381 | 380 | 1 | -3748.60 | -3748.60 | 0.00054 | **<u>0.01857</u>** |
| *ND3* | 381 | 380 | 1 | -867.15 | -867.15 | 0.00014 | **<u>0.00937</u>** |
| *ND4* | 381 | 380 | 1 | -3826.87 | -3826.87 | -0.00067 | 1.00000 |
| *ND4L* | 381 | 380 | 1 | -781.98 | -781.98 | 0.00000 | **<u>0.00113</u>** |
| *ND5* | 381 | 380 | 1 | -5019.63 | -5019.63 | 0.00173 | **<u>0.03318</u>** |
| *ND6* | 381 | 380 | 1 | -1931.80 | -1931.80 | -0.00013 | 1.00000 |

| *Bos indicus* mitochondrial genome sequences |  |  |  |  |  |  |  |
| --- | --- | --- | --- | --- | --- | --- | --- |
| Gene symbol | No. param. $H_A$ | No. param. $H_0$ | df | LnL $H_A$ | LnL $H_0$ | LRT($2\Delta\text{LnL}$) | $P$ ($\chi^2$) |
| *ATP6* | 381 | 380 | 1 | -1894.03 | -1894.03 | 0.00019 | **<u>0.01090</u>** |
| *ATP8* | 381 | 380 | 1 | -526.98 | -526.98 | 0.00013 | **<u>0.00900</u>** |
| *CYB* | 381 | 380 | 1 | -3334.59 | -3334.59 | 0.00193 | **<u>0.03510</u>** |
| *COX1* | 381 | 380 | 1 | -3955.67 | -3955.67 | 0.00021 | **<u>0.01170</u>** |
| *COX2* | 381 | 380 | 1 | -1696.06 | -1696.06 | -0.00057 | 1.00000 |
| *COX3* | 381 | 380 | 1 | -1774.40 | -1774.40 | 0.00104 | **<u>0.02570</u>** |
| *ND1* | 381 | 380 | 1 | -2475.83 | -2475.84 | 0.00159 | **<u>0.03190</u>** |
| *ND2* | 381 | 380 | 1 | -3747.84 | -3747.84 | -0.00085 | 1.00000 |
| *ND3* | 381 | 380 | 1 | -866.65 | -866.65 | -0.00031 | 1.00000 |
| *ND4* | 381 | 380 | 1 | -3826.87 | -3826.87 | -0.00053 | 1.00000 |
| *ND4L* | 381 | 380 | 1 | -781.98 | -781.98 | -0.00041 | 1.00000 |
| *ND5* | 381 | 380 | 1 | -5019.63 | -5019.63 | -0.00374 | 1.00000 |
| *ND6* | 381 | 380 | 1 | -1931.80 | -1931.80 | -0.00007 | 1.00000 |

| **Bos species group mitochondrial genome sequences** |  |  |  |  |  |  |  |
| --- | --- | --- | --- | --- | --- | --- | --- |
| **Gene symbol** | **No. param. $H_A$** | **No. param. $H_0$** | **df** | **LnL $H_A$** | **LnL $H_0$** | **LRT($2\Delta\text{LnL}$)** | **$P$ ($\chi^2$)** |
| *ATP6* | 381 | 380 | 1 | -1894.03 | -1894.03 | -0.00045 | 1.00000 |
| *ATP8* | 381 | 380 | 1 | -527.49 | -527.49 | -0.00043 | 1.00000 |
| *CYB* | 381 | 380 | 1 | -3334.59 | -3334.59 | 0.00042 | **<u>0.01639</u>** |
| *COX1* | 381 | 380 | 1 | -3955.67 | -3955.67 | -0.00001 | 1.00000 |
| *COX2* | 381 | 380 | 1 | -1696.06 | -1696.06 | 0.00000 | 1.00000 |
| *COX3* | 381 | 380 | 1 | -1774.40 | -1774.40 | -0.00017 | 1.00000 |
| *ND1* | 381 | 380 | 1 | -2475.83 | -2475.83 | 0.00027 | **<u>0.01311</u>** |
| *ND2* | 381 | 380 | 1 | -3748.60 | -3748.60 | 0.00026 | **<u>0.01291</u>** |
| *ND3* | 381 | 380 | 1 | -867.15 | -867.15 | -0.00137 | 1.00000 |
| *ND4* | 381 | 380 | 1 | -3825.86 | -3825.86 | -0.00035 | 1.00000 |
| *ND4L* | 381 | 380 | 1 | -781.98 | -781.98 | -0.00001 | 1.00000 |
| *ND5* | 381 | 380 | 1 | -5018.53 | -5018.53 | -0.00080 | 1.00000 |
| *ND6* | 381 | 380 | 1 | -1931.80 | -1931.80 | -0.00076 | 1.00000 |

**Table S4.** HMG and LMG functional subset genes detected in multiple studies (≥ 2) of genomic selective sweeps in African cattle populations.

| Gene symbol | Gene name | Functional subset | Chromosomal location | Citations |
| --- | --- | --- | --- | --- |
| *ACO2* | Aconitase 2 gene | LMG | BTA5 | (Bahbahani et al., 2018a; Taye et al., 2018) |
| *BCL2L13* | BCL2 like 13 gene | LMG | BTA5 | (Taye et al., 2018; Taye et al., 2017) |
| *CA5A* | Carbonic anhydrase 5A gene | LMG | BTA18 | (Kim et al., 2017; Taye et al., 2018; Taye et al., 2017) |
| *CMC1* | C-X9-C motif containing 1 gene | LMG | BTA22 | (Bahbahani et al., 2018a; Kim et al., 2017) |
| *CYP24A1* | Cytochrome P450, family 24, subfamily A, polypeptide 1 gene | LMG | BTA13 | (Bahbahani et al., 2017; Tijjani et al., 2019) |
| *DARS2* | Aspartyl-tRNA synthetase 2, mitochondrial gene | HMG | BTA16 | (Bahbahani et al., 2017; Taye et al., 2018) |
| *DNAJC11* | DnaJ heat shock protein family (Hsp40) member C11 gene | LMG | BTA16 | (Bahbahani et al., 2017; Taye et al., 2018; Taye et al., 2017) |
| *HDHD3* | Haloacid dehalogenase like hydrolase domain containing 3 gene | LMG | BTA8 | (Kim et al., 2017; Taye et al., 2017) |
| *HIBADH* | 3-hydroxyisobutyrate dehydrogenase gene | LMG | BTA4 | (Kim et al., 2017; Taye et al., 2018; Taye et al., 2017) |
| *HIGD1A* | HIG1 hypoxia inducible domain family member 1A gene | LMG | BTA22 | (Bahbahani et al., 2018a; Jang et al., 2021; Tijjani et al., 2019) |
| *MRPS33* | Mitochondrial ribosomal protein S33 gene | HMG | BTA4 | (Kim et al., 2017; Taye et al., 2017) |
| *MSRB3* | Methionine sulfoxide reductase B3 gene | LMG | BTA4 | (Bahbahani et al., 2017; Kim et al., 2017; Taye et al., 2018; Taye et al., 2017) |
| *NGRN* | Neugrin, neurite outgrowth associated gene | LMG | BTA21 | (Taye *et al.*, 2018; Taye *et al.*, 2017) |
| *PDE12* | Phosphodiesterase 12 gene | LMG | BTA22 | (Bahbahani *et al.*, 2017; Taye *et al.*, 2018) |
| *PUSL1* | Pseudouridine synthase like 1 gene | LMG | BTA16 | (Taye *et al.*, 2018; Taye *et al.*, 2017) |
| *SARDH* | Sarcosine dehydrogenase gene | LMG | BTA11 | (Bahbahani *et al.*, 2018a; Tijjani *et al.*, 2019) |
| *SLC25A11* | Solute carrier family 25 member 11 gene | LMG | BTA19 | (Bahbahani *et al.*, 2017; Kim *et al.*, 2017; Taye *et al.*, 2018) |
| *SLC25A21* | Solute carrier family 25 member 21 gene | LMG | BTA21 | (Bahbahani *et al.*, 2018b; Tijjani *et al.*, 2019) |
| *SOD1* | Superoxide dismutase 1 gene | LMG | BTA1 | (Kim *et al.*, 2017; Taye *et al.*, 2018; Taye *et al.*, 2017) |
| *SUCLG2* | Succinate-CoA ligase GDP-forming subunit beta gene | LMG | BTA22 | (Jang *et al.*, 2021; Kim *et al.*, 2020) |
| *TIMM21* | Translocase of inner mitochondrial membrane 21 gene | LMG | BTA24 | (Bahbahani *et al.*, 2018a; Tijjani *et al.*, 2019) |
| *TOMM22* | Translocase of outer mitochondrial membrane 22 gene | LMG | BTA5 | (Kim *et al.*, 2017; Taye *et al.*, 2017) |
| *YME1L1* | YME1 like 1 ATPase gene | LMG | BTA13 | (Taye *et al.*, 2018; Taye *et al.*, 2017) |

### Supplementary Information Table Captions

**Table S5:** *Bos* species mitochondrial genome sequences used for molecular evolution analysis.

**Table S6:** Functional gene subsets used to detect evidence for mitonuclear disequilibria in African admixed cattle populations.

